# The *Salix* SmSPR1 Involved in Light-Regulated Cell Expansion by Modulating Microtubule Arrangement

**DOI:** 10.1101/668889

**Authors:** Liu Xiaoxia, Zhang Jianguo, Sui Jinkai, Luo Ying, Rao Guodong

**Affiliations:** State Key Laboratory of Tree Genetics and Breeding, Research Institute of Forestry, Chinese Academy of Forestry, Beijing 100091, China; Collaborative Innovation Center of Sustainable Forestry in Southern China, Nanjing Forestry University, Nanjing 210037, China; Key Laboratory of Tree Breeding and Cultivation, National Forestry and Grassland Administration, Research Institute of Forestry, Chinese Academy of Forestry, Beijing 100091, China

**Keywords:** *Salix matsudana*, microtubule, microtubule-associated proteins, protein interaction, light regulation, cell expansion and elongation, SPR1

## Abstract

Light signaling and cortical microtubule (MT) arrays are essential to the anisotropic growth of plant cells. Microtubule-associated proteins (MAPs) function as regulators that mediate plant cell expansion or elongation by altering the arrangements of the MT arrays. However, current understanding of the molecular mechanism of MAPs in relation to light to regulate cell expansion or elongation is limited. Here, we show that the microtubule-associated protein SPR1 is involved in light-regulated directional cell expansion by modulating microtubule elongation in *Salix matsudana*. Overexpression of *SmSPR1* in Arabidopsis results in right-handed helical orientation of hypocotyls in dark-grown etiolated seedlings, whereas the phenotype of transgenic plants was indistinguishable from those of wild-type plants under light conditions. Phenotypic characterization of the transgenic plants showed reduced anisotropic growth and left-handed helical MT arrays in etiolated hypocotyl cells. Protein interaction assays revealed that SPR1, CSN5A (subunits of COP9 signalosome, a negative regulator of photomorphogenesis), and ELONGATED HYPOCOTYL 5 (HY5, a transcription factor that promotes photomorphogenesis) interacted with each other *in vivo*. The phenotype of Arabidopsis *AtSPR1*-overexpressing transgenic lines was similar to that of *SmSPR1*-overexpressing transgenic plants, and overexpression of *Salix SmSPR1* can rescue the *spr1* mutant phenotype, thereby revealing the function of SPR1 in plants.

**Highlight:** Function of microtubule-associated protein SPR1 is directly related to light, and crucial to the balance of tubulin polymerization

## Introduction

Plant cells exhibit different development patterns, ranging from skotomorphogenic development in darkness to photomorphogenesis. Cell proliferation of skotomorphogenesis in dark-grown seedlings does not significantly change, whereas the hypocotyl rapidly elongates in one direction, and this highly directional cell expansion results in the elongation of this specific organ. This process is accompanied by an apical hook, which is attenuated by pro-plastid differentiation that is essential to the initiation of a newly germinated seedling to push through the soil. In contrast, when seedlings are exposed to light, hypocotyl elongation is limited, petioles and cotyledons open, pro-plasmids develop into chloroplasts, and roots elongate (And and Deng, 1996; Gendreau *et al*., 1997). Previous studies have demonstrated that many factors mediate cell expansion and elongation in the dark and light, which include multiple photoreceptors, plant hormones, and transcription factors (Castillon *et al*., 2007; Galvão and Fankhauser, 2015). These studies have largely concerned with the identification of upstream influence factors in mediating dark- or light-related signal pathways, and how plants coordinate downstream regulators during cell expansion is still unclear.

Both genetics and physiology support the view that the arrangement of cortical microtubule (MT) participates in the regulation of cell expansion and elongation. In rapidly extending cells such as the tissues of the hypocotyl in etiolated seedlings, cortical MTs are predominantly arranged in parallel with each other, which results in transverse growth to the growth direction (Galva *et al*., 2014; Sedbrook and Kaloriti, 2008). In contrast, as cell elongation slows down, the arrangements of the MTs shift from parallel to oblique or longitudinal in direction (Barker *et al*., 2010; Crowell *et al*., 2011). These MT arrangements guide the positioning and trajectories of cellulose-synthesizing protein complexes as these track along cortical MTs beneath the plasma membrane and deposit cellulose microfibrils around the entire cell, and the orientation of microfibrils cross-lined with hemicelluloses mainly determines cell expansion and elongation (Brandizzi and Wasteneys, 2013; Foster *et al*., 2003; Fujita *et al*., 2012). The transverse orientation of cortical MTs in etiolated hypocotyl cells is reorganized into an oblique or longitudinal array when seedlings are exposed to light, which also slows down cell elongation in the hypocotyl (Sambade *et al*., 2012), indicating a connection between cortical MT arrangements and light signals.

The arrangement of MT arrays is related to plant morphology, and rearrangements of cortical MTs from transverse to left-handed (right-handed) helical or oblique alignment have been proposed to drive cells from elongation to expansion, which results in the twisted growth of plant organs (Galva *et al*., 2014). Microtubule-associated proteins (MAPs) regulate the organization and dynamics of MTs. Numerous studies have shown that MAP regulates cell expansion and elongation by altering the arrangement and dynamics of cortical MT (Lian *et al*., 2017; Lucas *et al*., 2011; Shoji *et al*., 2004; Sun *et al*., 2015). SPIRAL1 (SPR1) is a plant-specific MAP that was identified in an *Arabidopsis spr1* mutant that exhibited helical root growth. When grown on a tilted hard-agar surface, right hand spiral appears in the *spr1* epidermal cells of the root, and the roots of *spr1* exhibit right directional growth when viewed from above the agar plates(Furutani *et al*., 2000). The arrangement of cortical MTs in root elongation zone showed a phenotype of the left-handed helix in the *spr1* mutant, rather than parallel alignment in the wild-type plants (Nakajima *et al*., 2004). From these observations, Furutani et al. therefore concluded that SPR1 plays an important role in maintaining the function integrity of cortical MTs and is essential for anisotropic expansion of cells. Previous studies have shown that SPR1 binds to another plus-end tracking protein EB and synergistically regulates the polymerization and elongation of MTs (Furutani *et al*., 2000; Galva *et al*., 2014). SPR1 can also be ubiquitinated under salt stress by 26S proteasome and accelerate the depolymerization and reorganization of MTs, which is required for plant salt stress tolerance (Wang *et al*., 2011). Whether SPR1 has other interacting proteins that are involved in regulating polymerization and elongation of MTs remains unclear.

*Salix* (willow) are widely distributed, from North America to China, and contain more than 300 species varying from small shrubs to trees(Barker *et al*., 2010). The phenotypes of *Salix matsudana* and its varieties are diverse, including the phenotype of branch spiral, the phenotype of vertical growth of branch, and the spherical phenotype of crown, making it a good material for studying tree morphology. In this study, we identified six *SmSPR1* genes from *S. matsudana*, and qRT-PCR-based tissue-specific transcript abundance analysis showed that *SmSPR1* is upregulated in all tissues tested. We then analyzed this gene in greater detail. Overexpression of *SmSPR1* in *Arabidopsis* resulted in hypocotyl helical growth in etiolated seedlings, whereas no hypocotyl helical growth or root twisting was observed in transgenic seedlings growth in the presence of light. To rule out that the helix phenotype was caused by heterologous expression, we overexpressed the *Arabidopsis AtSPR1* gene and obtained the same phenotype. We then transferred the helical etiolated seedling to light conditions, which resulted in straight new upper hypocotyls, and the formed lower stem also showed the right-handed helical orientation. However, the straight light-grown hypocotyl twisted when the transferred to the dark. Given that light regulates the helical growth of seedling hypocotyls, we then set out to identify light-related proteins that interact with SmSPR1. Here, we show that SPR1, CSN5A, and HY5 interacted with each other *in vivo*, and influencing anisotropic cell growth in *Arabidopsis*. We propose a model that SPR1 mediates the strength balance of MTs, either via loss of function or overexpression of *SPR1*, eventually resulting in the helical growth of MTs.

## Materials and Methods

### Plant materials

*S. matsudana* Koidzin in this study were collected from the Beijing Botanical Garden. The leaves, annual shoot tips, and stems were frozen in liquid nitrogen and then stored at −80°C. All *A. thaliana* plants were of the Columbia-0 ecotype (Col-0). Mutant *spr1* seeds (CS6547) and 35S:Tubulin6B-GFP (CS6550) transgenic seeds were obtained from the Arabidopsis Biological Resource Center (ABRC; https://www.arabidopsis.org/).

For plant physiological analysis, the seeds were surface-sterilized for 1 min with 70% (v/v) ethanol and then washed with 15% (v/v) sodium hypochlorite (∼10%) for 12 min. The transformed seeds were sown on MS plates with 3% sucrose and 0.6% agar containing 50 mg/L kanamycin for mutant selection or 25 mg/L phosphinothricin for transgenic selection. For phenotypic analysis and biochemical assays, the seeds were placed on half-strength MS medium containing 0.8% agar and 1% sucrose. For the hypocotyl measurements, the plates were transferred to 22°C in the light or dark for 7 d after stratification at 4°C for 3 d.

### Sequence identification and analysis of SmSPR1 promoter

*Populus* and Arabidopsis *CSN5A, COP1, HY5*, and *SPR1* family genes were used for identifying the CDS homologs of the *SPR1, CSN5A, COP1*, and *HY5* genes in *S. matsudana* using BLAST. The primers used for amplification of full-length *SPR1, CSN5A, COP1* and *HY5* are listed in Supplemental Table S1. Phylogenetic analysis was performed using the software MEGA 6(Tamura *et al*., 2013). The phylogenetic relationship of the genetic model was assessed using neighbor-joining tree with 1,000 bootstrap trials.

The promoter of *SmSPR1* was subjected to homology-based cloning according to the genome sequence of *Salix suchowensis* and *Populus trichocarpa*. The sequence and then cloned into *pBI121* vector, replacing the *Cauliflower mosaic virus* (CaMV) 35S promoter, to drive the *GUS* (β-glucuronidase) reporter gene. Then the recombinant *pBI121*-*SmSPR1*::*GUS* construct was transformed into *Tabaco* plants using *Agrobacterium tumefaciens* (strain GV3101). The stem, root, and petiole section of transformed plantlets and the 10-days-old seedlings were used for histochemical staining. The GUS staining procedure was performed as described according to previous study (Hwang *et al*., 2014).

### Semi RT-PCR and real-time PCR

Semi RT-PCR was used for determination the expression level of *SmSPR1* and *AtSPR1* in the wild-type and transgenic plants. Arabidopsis*18S rRNA* (At3G41768) was used as a loading control. The primers used for these assays are described in Supplemental Table S2.

Real-time PCR was performed for the quantification of *SmSPR1* family member transcripts in the tissue of shoot tips, roots, stems, xylem, and phloem. The RNA was isolated from *S. matsudana* using Plant RNA extraction kit (CWbio. Co., Ltd.) and SYBR GreenI Taq Mix (CWbio. Co., Ltd.) was used as fluorochrome. The *SmSPR1* gene family primers are listed in Supplementary Table S3. GAPDH was used as internal standard. All reactions were repeated at least three times under identical conditions.

### Generation of SmSPR1 and AtSPR1 overexpression transgenic plants

For the *spr1* mutant of the Arabidopsis complement test and *AtSPR1* overexpression assay, *SmSPR1* and *AtSPR1* cDNA was amplified and introduced into pDONR221 via BP reaction and to pEarleyGate104 of Gateway vectors via LR recombinase (Invitrogen) (Earley *et al*., 2010). All primers are listed in Supplemental Table S4. The resulting constructs were transformed into Arabidopsis using *Agrobacterium tumefaciens* (GV3101) via Arabidopsis floral dip method as described elsewhere (Zhang *et al*., 2006). The homozygous T3 seedlings were used for further analyses.

### Phenotypic analysis

The spiral phenotypes of seedlings were observed using an Ultra depth of field microscope (Leica DVM6) equipped with CCD (PLANAPO FOV 12.55). For measuring length and width of hypocotyl and root, the relevant parameter was measured using ImageJ (http://rsb.info.nih.gov/ij/). Hypocotyls of five-day-old seedlings were fixed with 50% FAA and then embedded in spr resin (SPI). A series of 4-μm thick longitudinal sections and transverse sections (rapidly elongating region) were made with a rotary microtome RM2265 (LEICA EM UC7). Fixed sections were stained with toluidine blue O for 30 min and photographed with an Olympus BX51 microscope equipped with a DP74 camera.

For drug treatment, wild and transgenic seeds were grown on 0.8% agar-solidified 1/2 MS vertically oriented plates for 7 days with or without specific concentration of Propyzamide (Sigma-Aldrich). To detect the morphology of the cells of 7-day-old seedlings, etiolated hypocotyls and roots were soaked in 10 μM PI (Sigma-Aldrich). Zeiss LSM510 confocal microscope (with 543 nm diode laser, and an emission band of 560 to 690 nm) was used for images collecting.

For measurement of MT arrays, *SmSPR1* was over-expressed in *35S: GFP-TUB6* background and detected on a Zeiss LSM510 confocal microscope. The orientation of cortical MTs in epidermal cell was measured at upper regions. Measurements were performed using ImageJ (http://rsb.info.nih.gov/ij/). Microtubules with clear visible were selected for measurements in each cell (n ≥ 25cells). The procedure was performed as previously described (Liu *et al*., 2013).

### Protein Purification and Pull-down assay

The pET28a-SmSPR1-His (SmSPR1-His), pET21a-HY5-Flag (HY5-Flag) and pGEX4T1-GST-SmCSN5A-Flag (GST-SmCSN5A-Flag) clones were transformed into the Rosetta. All primers for constructing prokaryotic expression vectors are listed in Supplemental Table S5. The protein prokaryotic expression and purification was according to the instruction of Ni-NTA Agarose (Cat.No.30210, QIAGEN) and Glutathione Sepharose (GE).

### Yeast Two-Hybrid analysis

The CDS of *SmCSN5A, SmCOP1* and *SmHY5* were constructed on the yeast two-hybrid prey vector pGADT7, respectively. SmSPR1 was constructed on the bait vector pGBKT7. The primers were listed in Supplemental Table S6. The bait vector of *SmSPR1* was transformed with *SmCSN5A, SmCOP1* and *SmHY5* respectively into the yeast strain AH109 as instructions for Matchmaker GAL4 Two-Hybrid Systems 3 (Clontech). Yeast Two-Hybrid analysis were performed following Yeast Protocols Handbook (Clontech). Transformed yeast cells were separately spread to 2D synthetic deficiency medium (SD-TL: -Trp/-Leu) and 4D selective medium (SD-TLHA: -Trp/-Leu/-His/-Ade) and then placed at 30°C for 4 days. Saturated yeast cells were dilute to 1:1, 1:10, 1:100, 1:200, 1:500, and 1:1,000, and then spotted onto the selection medium.

### Bimolecular Fluorescence Complementation (BiFC) Assay

BiFC assays were performed using pEarleyGate vectors (pEarleyGate201-YN, pEarlyGate202-YC). The vectors were kindly supplied by Dr. Bin Tan (Yi *et al*., 2013). The cDNA encoding SmCSN5A, SmCOP1, and SmHY5 were fused with the C-terminal fragment of YFP. SmSPR1 was fused to the fragment encoding the N-terminus of YFP. The primers used for BiFC were listed in Supplemental Table S7. All vectors were transformed into Agrobacterium strain (GV3101). The transient expression was according to the method of (Sparkes *et al*., 2006). Detection of BiFC signals was visualized by Zeiss LSM510 microscope. The excitation light had a wavelength of 514 nm, and the emission light receiving range was 525-555 nm. The experimental results were analyzed using Zen software.

### Statistical analyses

The quantitative data were analyzed using a one-variable general linear model procedure (ANOVA) with the SPSS software package (SPSS Inc., http://www.spss.com.cn). Analysis of significance was performed using t test or Duncan’s multiple range tests at P≤0.05 or P≤0.01. Data are reported as the mean ± SE of three or more experiments.

### Accession numbers

Sequence data from this article can be found in the GenBank data libraries under accession numbers: *SmSPR1*, MK770432; *SmSPR1_L1*, MK770433; *SmSPR1_L2*, MK770434; *SmSPR1_L3*, MK770435; *SmSPR1_L4*, MK770436; *SmSPR1_L5*, MK770437; *SmCOP1*, MK770438; *SmCSN5A*, MK770439; *SmHY5*, MK770440.

## Results

### Cloning and Expression Pattern of the *SmSPR1* Genes

A total of six *SmSPR1* genes were identified according to the sequences of the Arabidopsis *SPR1* family genes (Nakajima *et al*., 2006; Sedbrook *et al*., 2004). These *SmSPR1s* were then isolated from *S. matsudana* using PCR-based approaches with gene-specific primers (Supplemental Table S1). We named these *Salix SPR1* genes as *SmSPR1* and *SmSPR1-LIKE* genes (*SmSPR1_L1-SmSPR1_L5*) based on their amino acids sequence identity and phylogenetic relationship with Arabidopsis *SPR1* family genes (Fig. 1A). All *Salix* and Arabidopsis SPR1s were classified into three classes (designated to Class I-III): Class I included *Salix* SPR1, SPR1_L3, SPR1_L4, with Arabidopsis SPR1 and SPRL2; Class II consisted of *Salix* SPR1_L1 and SPR1_L2 and Arabidopsis SPR1L3, SPR1L4 and SPR1L5; and Class III comprised *Salix* SPR1_L5. *Salix* SPR1 and its homolog sequences shared N- and C-terminal regions except SmSPR1_L5, and the outgroup position of SmSPR1_L5 may be due to the absent of conserved C-terminal region (Fig. 1B). Highly conserved repeat amino acids sequences were observed at the N- and C-termini in SmSPR1, L1, L2, and L4, with the consensus motif being GGG/DQ/SSSLG/DY/FLFG (Fig. 1B). At the C-terminal of this conserved motif, the PGGG sequence is present in many mammalian MAPs and is a conserved binding sequence of microtubules (Nakajima *et al*., 2004).

**Fig. 1.**
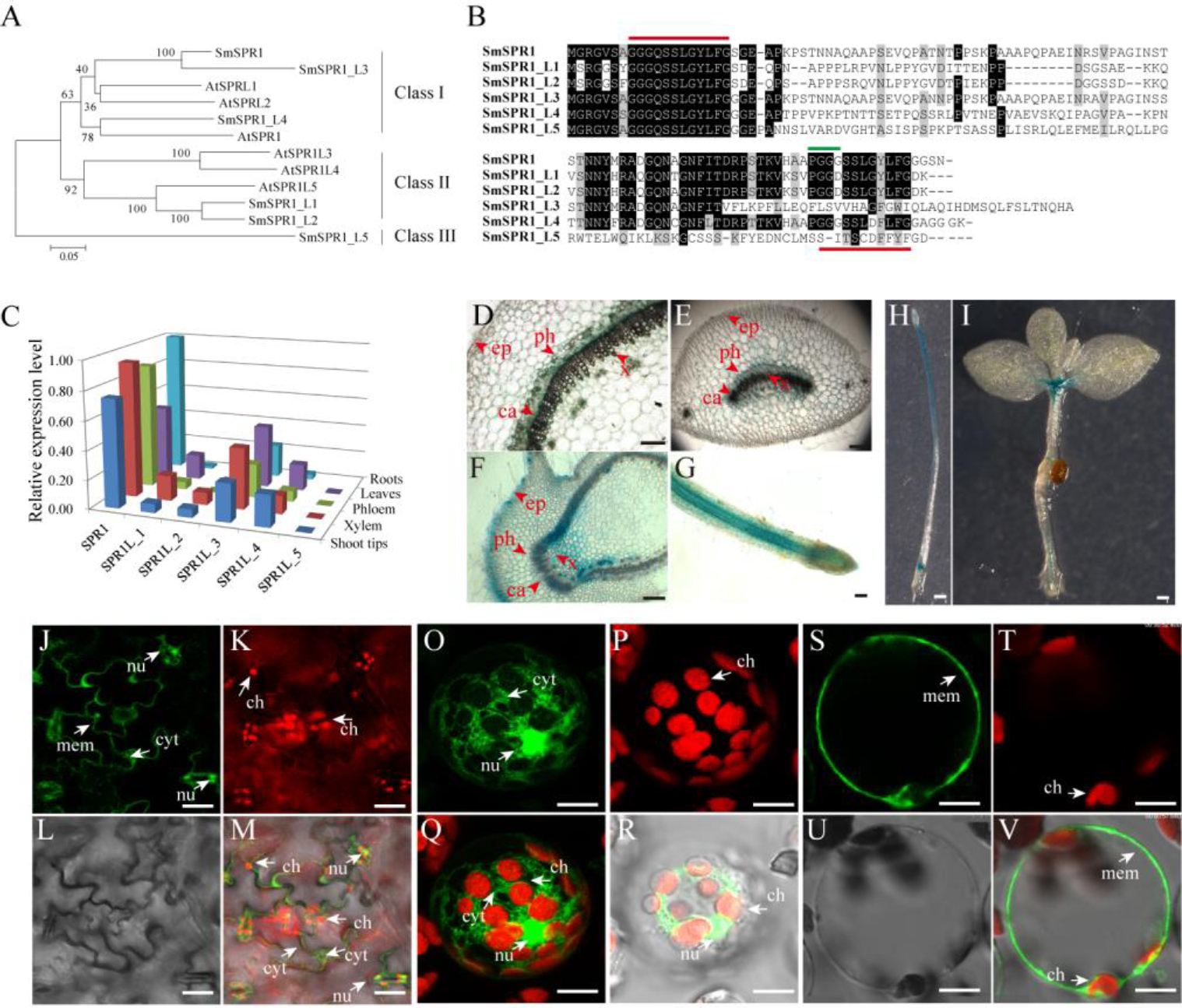
Gene sequence analysis, tissue-specific expression, and cellular localization of SmSPR1. (A) An unrooted phylogenetic tree was generated using full-length protein sequences of *Salix* and *Arabidopsis* SPR1 family isoforms. A neighbor-joining phylogenetic tree was generated using MEGA 6.0 with 1,000 bootstrap replicates. (B) Protein sequence alignment of *Salix* SPR1. Conserved residues are shown in black boxes. ClustalW with default settings was used for protein alignment. (C) qRT-PCR-based transcript abundance of six *Salix SPR1* genes. Expression levels were normalized to the geometric mean of three housekeeping genes (*GAPDH*). (D) to (I) Histochemical assay for GUS activity in transgenic tobacco lines. D-G, Transversal section of the of stem, midvein, axillary bud; f, Root; H, seedling grown in the dark for 12 days; I, Twenty-day-old seedling grown under light. Bars in (D-I) = 100 µm. (J) to (V) Subcellular localization of SmSPR1 in Arabidopsis epidermal cells and mesophyll protoplasts.J, O and S are fluorescence signals of GFP; K, P and T are fluorescent in the chloroplast; L and U under brightfield optics; m, r, and v are merged by GFP, chloroplast and brightfield; Q is merged with GFP and chloroplast. Ca, cambium; x, xylem; ph, phloem; ep, epidermis; nu, nucleus; ch, chloroplast; mem, cytomembrane; cyt, cytoplasm; bars in (J-V) = 10 µm.

To determine *SmSPR1* expression level at different tissues, we conducted a qRT-PCR-based tissue-specific transcript abundance analysis of five tissues (shoot tips, xylem, phloem, leaves, and roots) from three trees with gene-specific primers, and the transcript expression level of six *Salix SPR1* genes are shown in Fig. 1C. Class I *SPR1* gene had the highest expression level and was detected at almost equal levels in all tissues tested. *SPR1_L3* and *SPR1_L4*, which also belong to Class I, were expressed in all tissues, with a moderate transcript level compared to *SPR1*. The expression levels of Class II *SPR1_L1* and *SPR1_L 2* were significantly lower than those of the Class I genes, and extremely low transcript levels were observed in the roots of *SPR1_L2* and *SPR1_L5*. The relative expression levels of Class I *SPR1* members were higher than those of Class II and III *SPR1* members, suggesting the Class I *SPR1* family genes, especially *SmSPR1*, is the major gene in *Salix*, similar to the results of Arabidopsis *AtSPR1*, which also had a predominant transcript level in all tissues tested (Nakajima *et al*., 2004).

To further obtain details on the tissue specific expression pattern of *SmSPR1*, we generated transgenic tobaccos (*Nicotiana tabacum*) of *P*_*SmSPR1*_: *GUS* which were then stained for GUS for GUS activity test. Strong GUS activity was observed at the internodes of stems, including phloem, cambium, and xylem, but not in epidermal cells (Fig. 1D). Midveins at each internode were also stained for GUS activity, and a similar expression pattern was obtained as that observed in the stems; GUS staining was also observed in vascular tissues, which included strong GUS staining in the phloem, moderate GUS staining of the cambium and xylem, and negative GUS staining of the epidermal cells (Fig. 1E). Axillary buds and root tips showed significantly strong GUS activity in all tissues tested (Figs. 1F, G), indicating that *SmSPR1* has a high transcript expression level in the meristem and elongation zone. Transgenic tobacco seedlings, which were grown both in the dark and under continuous light, were also stained for the GUS activity. Strong GUS activity was detected in the hypocotyls and roots of dark-grown seedlings (Fig. 1H), and high *SmSPR1* expression levels were detected in the shoot tips and roots of seedlings grown in light conditions (Fig. 1I).

To investigate the subcellular localization of *SmSPR1*, A *P35S-SmSPR1-GFP* fusion construct were generated and transiently expressed in Arabidopsis using a *A. tumefaciens*-mediated transformation approach. In non-plasmolyzed Arabidopsis leaf epidermal cells, the SmSPR1-GFP fusion protein was detected in the cell periphery, nuclei, and cytoplasm, but not in chloroplasts (Fig. 1J-M). The subcellular localization of SmSPR1 was examined by expressing the SmSPR1-GFP fusion protein in protoplasts prepared from Arabidopsis suspension-cultured cells, which was observed as a strong fluorescence signal, and the SmSPR1-GFP fusion protein was also observed in the cell periphery, nuclei, and cytoplasm, but not in chloroplasts (Fig. 1O-V).

### *SmSPR1* Transgenic Seedling Phenotype

To determine the function of SmSPR1, the *P35S*: *SmSPR1* transgenic Arabidopsis in the wild-type Col-0 background were generated. We identified 14 *P35S*:*SmSPR1* transgenic lines, all of which showed similar phenotypes. The phenotype of the transgenic plants was indistinguishable from those of the wild-type plants in the presence of light (Supplemental Fig. S1). This result also coincided with the phenotype of Arabidopsis *SPR1* overexpression transgenic lines (Nakajima *et al*., 2004). However, the hypocotyls in dark-grown seedling exhibited a right-handed helical orientation to the epidermal cells (Figs. 2A-B, Supplemental Fig. S2).

**Fig. 2.**
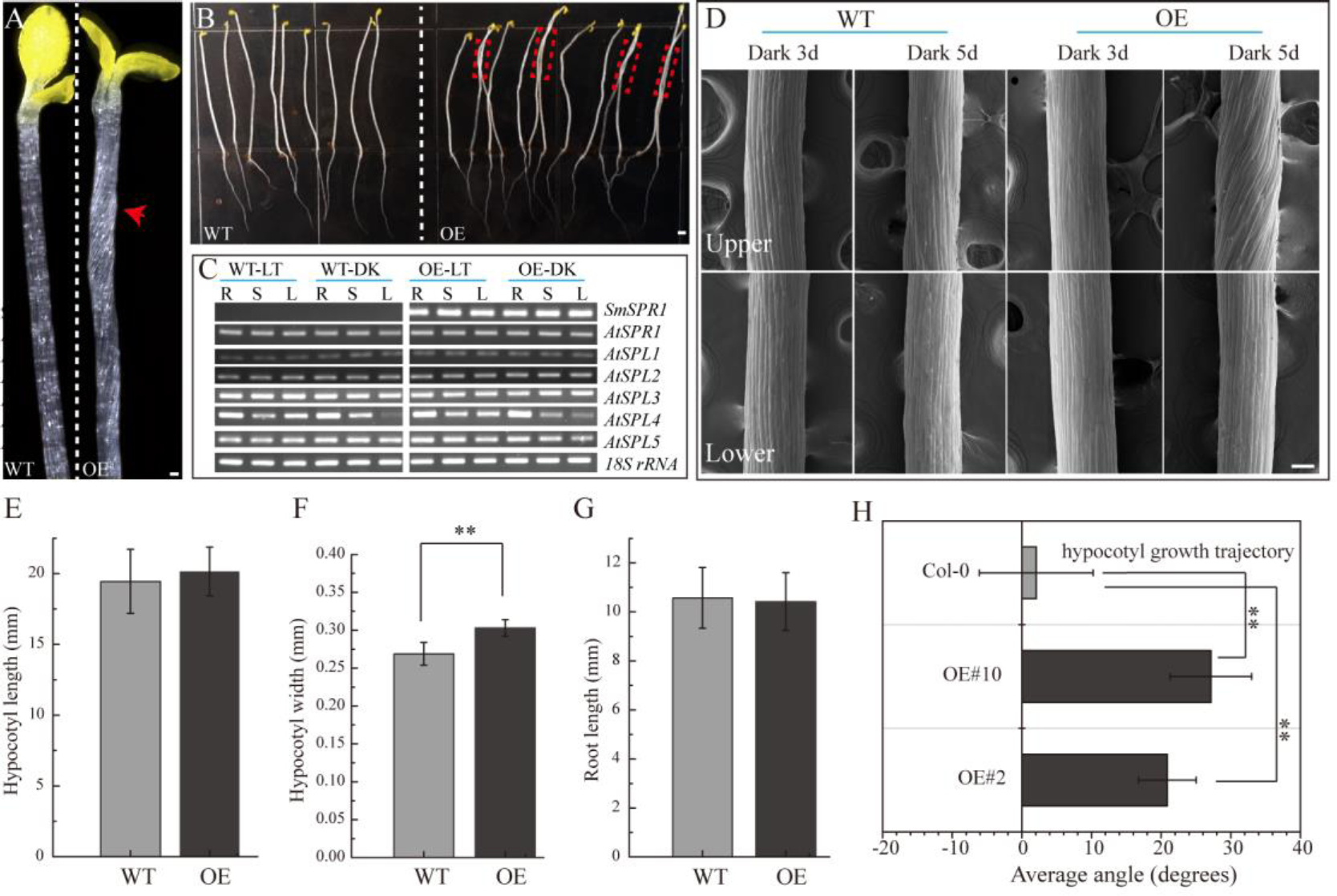
Phenotypes of transgenic plants. (A) *P35S: SmSPR1* transgenic plant showing a right-handed helical hypocotyl in etiolated seedlings compared to the wild-type. (B) Hypocotyl of *P35S: SmSPR1* transgenic plants showed right-tilted growth compared to the wild-type. (C) Semi-quantitative RT-PCR analyses of *SmSPR1* and *AtSPR1* genes in the roots (R), stems (S), and leaves (L). (D) Scanning electron microscopy of the upper and lower hypocotyls of etiolated transgenic seedlings. (E) Hypocotyl length of etiolated seedlings of transgenic and wild-type plants. (F) Hypocotyl width of etiolated seedlings in transgenic and wild-type plants. (G) Root length of etiolated seedlings of transgenic and wild-type plants. (H) Hypocotyl growth trajectory of etiolated seedlings of transgenic and wild-type plants (seven-day old seedlings).For (E) to (H), data are expressed as the mean ± SD of >30 seedlings. Asterisks indicate significant differences using the Student’s *t* test (*P* < 0.01). Bars in (A) to (D) = 100 µm.

To test whether the right-handed helical phenotype is the result of posttranscriptional gene silencing, semi-quantitative RT-PCR analysis was performed to determine the transcription levels of the *SmSPR1* and Arabidopsis *SPR1* family genes. A total of six Arabidopsis *SPR1* family genes were tested, and each gene showed moderate expression levels both in the wild-type and transgenic plants. High levels of *SmSPR1* transcripts were observed in the *P35S*:*SmSPR1* transgenic plants, whereas no *SmSPR1* transcripts were detected in the wild-type plants that were grown separately in dark and light conditions (Fig. 2C). These results suggested that the right-handed helical phenotype could be attributed to the high transcription levels of *SmSPR1* rather than Arabidopsis *SPR1s* posttranscriptional gene silencing. In addition, this right-handed helical phenotype was only observed after three days of growth in the dark, and it is notable that only the upper hypocotyl of etiolated seedlings showed the right-handed helical orientation (Fig. 2D, Supplemental Figs. S2 and S3). No differences in hypocotyl and root length were observed, although hypocotyl width significantly differed between the transgenic and wild-type plants (Student’s *t* test, P < 0.01, Figs. 2E-H). Considerable evidence indicates that Arabidopsis *spr1* mutants exhibit root morphological changes, which include twisted root epidermal cells and right directional root development when grown on vertically oriented hard agar plates (Nakajima *et al*., 2004; Nakajima *et al*., 2006; Sedbrook *et al*., 2004). We then surveyed the appearance of roots in the *P35S*:*SmSPR1* transgenic and wild-type plants, and the phenotypes of the roots were also indistinguishable from those of the wild type that were grown in the dark (Supplemental Fig. S2).

To investigate changes in the phenotype of *SmSPR1* overexpression plants at the cellular level, resin-embedded transverse and longitudinal sections of the upper region of etiolated hypocotyls of the seedlings were prepared. The results showed that the cross-sectional hypocotyl area of the transgenic plants was significantly larger than that of wild-type plants, and this enlargement was caused by the expansion of cells (Figs. 3A, B). The longitudinal section also showed bigger hypocotyls as well as cell expansion (Figs. 3C, D). Next, the *SmSPR1* gene was overexpressed in the *P35S:GFP:AtTUA6* Arabidopsis background. We then observed the arrangement of MTs of etiolated hypocotyls in the wild-type and transgenic plants. No differences in the MTs were observed between the wild-type and transgenic plants that were grown in the presence of light (Figs. 3E, F). In etiolated seedlings, the arrangement of MTs in the hypocotyl of wild-type plants was mainly parallel to each other and perpendicular to the long axis, whereas the MT arrays of the transgenic plants predominantly showed left-handed spiral growth (Figs. 3G, I).

**Fig. 3.**
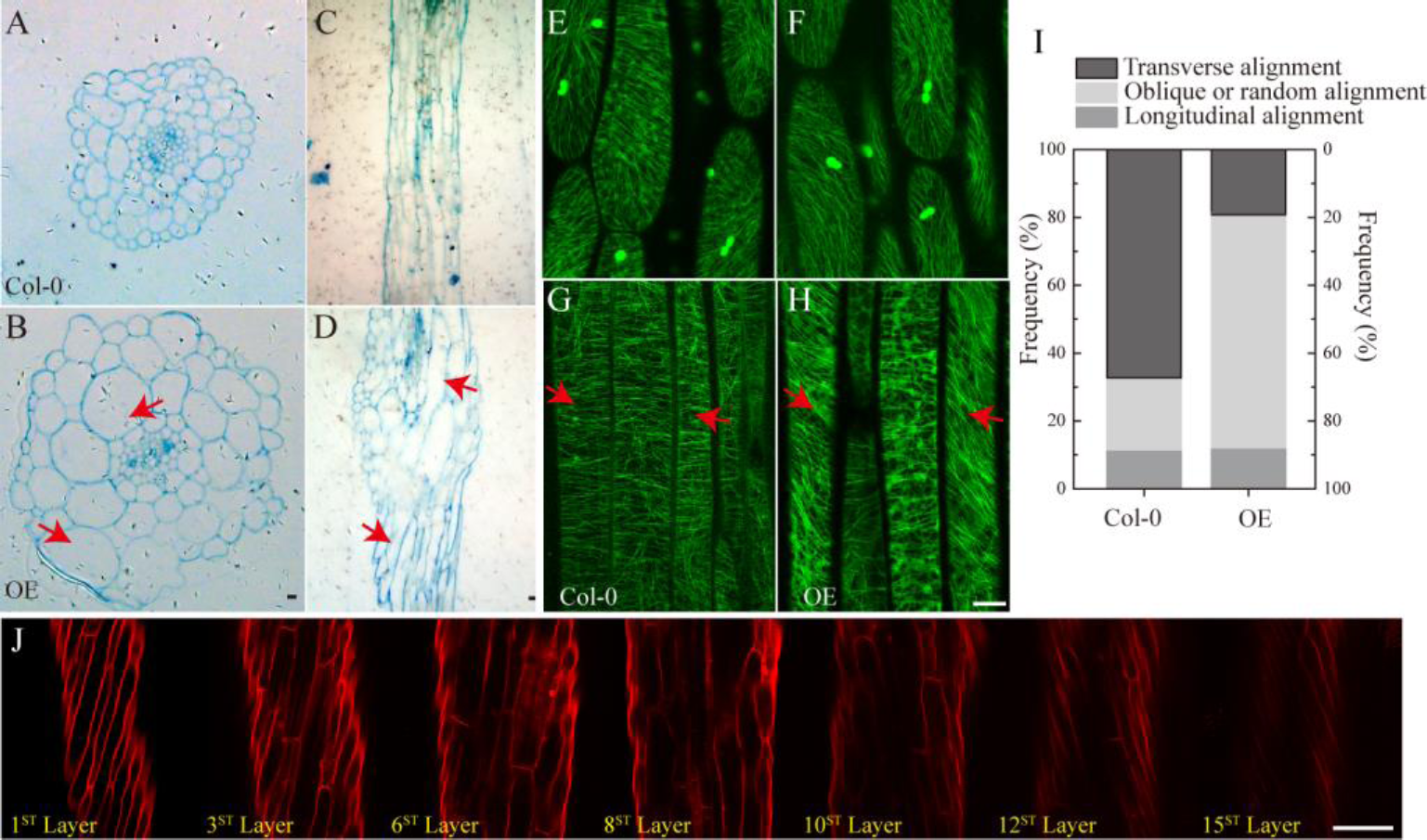
Cellular and MT differences between wild-type and transgenic plant. (A) Cross-sectional hypocotyl area of the wild-type plants. (B) Cross-sectional hypocotyl area of the transgenic plants. (C) Longitudinal section hypocotyl area of the wild-type plants. (D) Longitudinal section hypocotyl area of the transgenic plants. (E) MT arrangement in the hypocotyl of wild-type Arabidopsis grown in the presence of light. (F) MT arrangement in the hypocotyl of transgenic Arabidopsis grown in the presence of light. (G) MT arrangement in the hypocotyl of wild-type Arabidopsis grown in the dark. (H) MT arrangement in the hypocotyl of transgenic Arabidopsis grown in the dark. (I) Frequency of different MT orientation patterns in dark-grown hypocotyls of wild-type and transgenic plants. The data are expressed as the mean ± SD of > 30 seedlings. (J) Cell shapes and sizes of different layers from one side to the opposite side of hypocotyls in the transgenic etiolated seedling. Bars in (A) to (J) = 100 µm.

Taken together with the fact that transgenic seedling epidermal cells had right-handed helical orientation in the absence of light, that finding raised the question of whether the internal region is also affected by the overexpressed *SmSPR1* gene. To assess this, confocal microscopy was performed assess cell shapes and sizes layer by layer, and a total of 15 cell layers was obtained from one side to the opposite side of the tissues. The angles of the right-handed helical orientation from the first layer to the middle layer (8th layer) gradually decreased. Scanning from the 8^th^ layer to 15^th^ layer revealed that the angle of helical orientation negatively increased to the left-handed helical orientation (Fig. 3J). In sum, the helical phenotype was observed in most layers of the hypocotyl and was not limited to the epidermal cells under dark condition. We also stained the tissues of seedlings grown in the presence of light with PI, all of which were indistinguishable between the wild-type and the transgenic plants (Supplemental Fig. 1D).

### Overexpression of SmSPR1 Results in Strong Tolerance to an MT-depolymerizing Drug

To localize the SmSPR1 protein, we constructed an SmSPR1:GFP fusion protein vector and transfected it into Arabidopsis. GFP signals were assessed by confocal laser scanning microscopy, which revealed filamentous structures in the hypocotyl, roots, leaves, and other tissues. Further observations showed that the green fluorescence signals of SmSPR1-GFP coincides with the immunofluorescence (red) of tubulin, demonstrating that SmSPR1 co-localizes with the MTs (Figs. 4A, Supplemental Fig. S4). A microtubule-depolymerizing drug, propyzamide (PPM), was then used to assess changes in *SmSPR1:GFP* transgenic lines. With the addition of PPM, no fluorescence signals of SmSPR1-GFP were observed, and after the removal of PPM, the green fluorescence of SmSPR1-GFP was again detected (Fig. 4B). This coincides with the depolymerization and reorganization of MTs, which also demonstrates the co-localization of SmSPR1 with MTs.

**Fig. 4.**
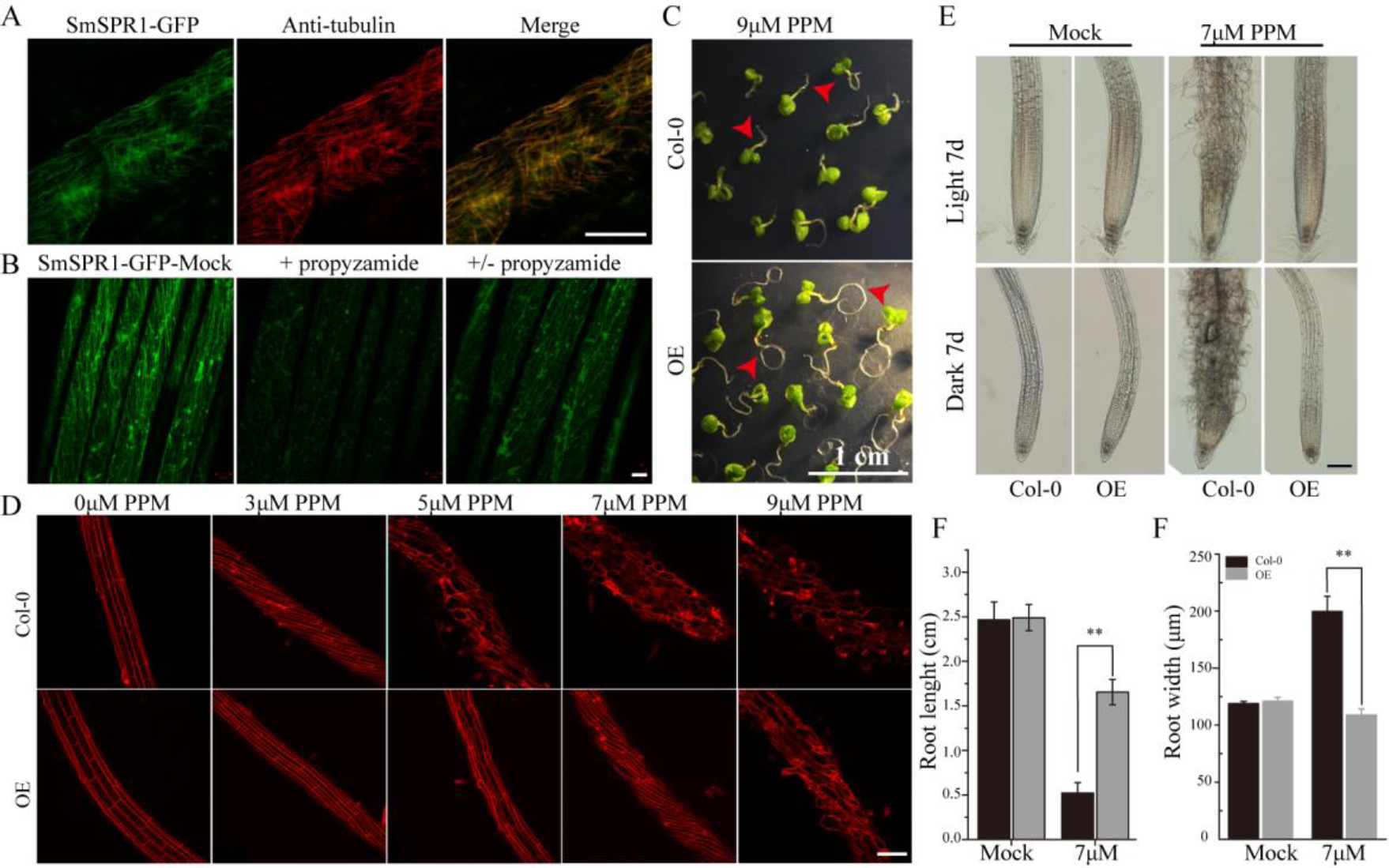
SmSPR1 protein localization and increase in PPM tolerance. (A) Confocal images of SmSPR1: GFP hypocotyl (green) and immunofluorescence stained with anti-tubulin antibodies (red). (B) SmSPR1: Changes in GFP fluorescence after the addition and removal of the MT-depolymerizing drug, PPM. (C) Seedling phenotypes of the wild-type and transgenic plants grown in a medium containing 9 µM PPM. (D) PI staining of roots of the wild-type and transgenic plants with higher concentrations of PPM in the presence of light. (E) Photomicrographs of roots from the wild-type and transgenic seedlings treated with 7 µM PPM. (F) Root length of wild-type and transgenic seedlings treated with 7 µM PPM. (G) Root width of the wild-type and transgenic seedlings treated with 7 µM PPM. For (F) to (G), the data are expressed as the mean ± SD of > 30 seedlings. Asterisks indicate significant differences using the Student’s *t* test (*P* < 0.01). Bars in (A) and (B) = 10 µm, (C) = 1 cm, (D) and (E) = 100 µm.

A previous study showed that the Arabidopsis *P35S:AtSPR1* line has a moderately higher resistance to long-tern treatment with PPM, and the maximum survival concentration of *P35S:AtSPR1* line is 5 µM (Nakajima *et al*., 2004). To determine whether SmSPR1 plays a similar function, *P35S:SmSPR1* transgenic seedlings were planted in agar medium containing PPM with different concentrations (Fig. 4, Supplemental Fig. S5). The roots of *P35S:SmSPR1* transgenic seedlings retained the ability to elongate compared to the wild type in the presence of 9 µM of PPM (Fig. 4C). PI staining showed that the roots of the wild-type plants exhibited a left-handed helical orientation with 3 µM of PPM, and the cells expanded or even dissociated at 5 µM, whereas roots of the transgenic plants showed a left-handed helical structure with 7 µM PPM, and the cells swelled at 9 µM (Figs. 4D-G). Our results indicated that the *P35S:SmSPR1* lines were highly tolerant of the MT-depolymerizing drug, surviving at a PPM concentration of 9 µM.

### Salix SmSPR1 and Arabidopsis AtSPR1 have Similar Biological Functions

To determine whether SmSPR1 has a similar biological function to Arabidopsis AtSPR1, we constructed an overexpression vector carrying the CaMV 35S promoter linked to *SmSPR1*, and transformed it into the Arabidopsis *spr1* mutant background. A total of 25 independent transgenic lines were generated, and their phenotypes were all analyzed in both light and dark conditions, and all transgenic lines showed similar phenotypes. The phenotype of the light-grown Arabidopsis *spr1* showed roots skewing to the right, which was rescued in the overexpression *SmSPR1* lines (Fig. 5A). In the dark, Arabidopsis *spr1* exhibited hypocotyl phenotype of right-handed helix, which was also rescued in the overexpression *SmSPR1* lines (Fig. 5B). Semi-quantitative RT-PCR analyses were used to assess the transcription levels of the *SmSPR1* gene in the wild, *spr1* mutant, and overexpression *SmSPR1* lines, which showed that only the *SmSPR1* overexpression lines expressed the *SmSPR1* gene (Fig. 5C). These results demonstrate that SmSPR1 and AtSPR1 have similar biological functions and play a role in maintaining anisotropic expansion of cells and straight growth of plants.

**Fig. 5.**
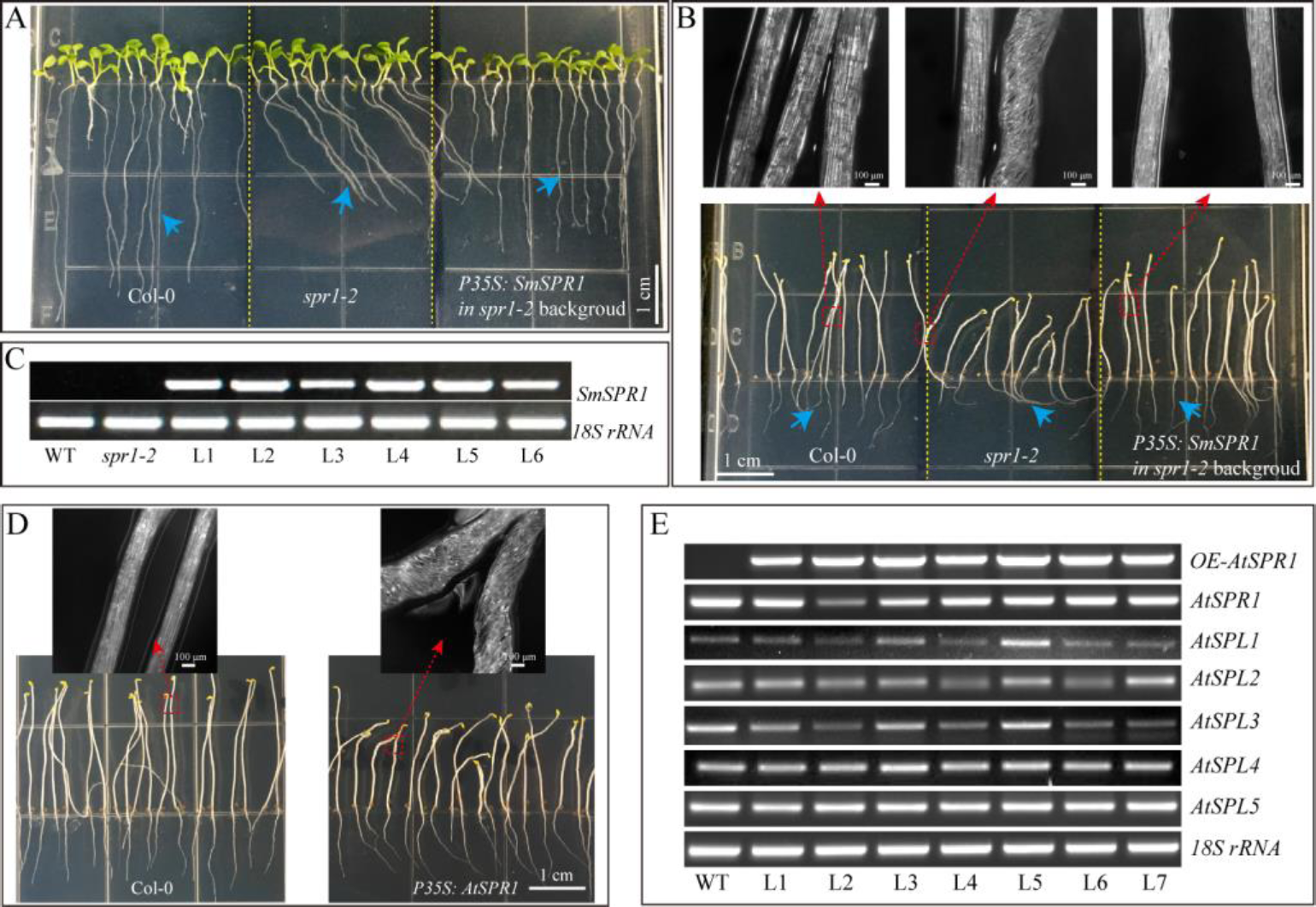
Overexpression of *SmSPR1* rescues in the *spr1* background and overexpression of Arabidopsis SPR1 causes a similar hypocotyl helix phenotype. (A) Phenotype of the wild-type, *spr1* mutant and overexpression *SmSPR1* transgenic plants in the *spr1* mutant background in the presence of light. (B) Etiolated seedling phenotype of the wild-type, *spr1* mutant and overexpression *SmSPR1* transgenic plants in the *spr1* mutant background; Blue short arrows show different root morphologies; Red boxes and long arrows indicate magnified hypocotyls. (C) Semi-quantitative RT-PCR analyses of the wild-type, *spr1* mutant and overexpression *SmSPR1* lines (L1-L6). (D) *P35S: AtSPR1* transgenic plant shows a right-handed helix hypocotyl in etiolated seedlings compared to the wild type. (E) Semi-quantitative RT-PCR analyses of wild type and overexpression *AtSPR1* lines (L1-L7).

To examine the generality of the SPR1 function in plants, we reconstructed the Arabidopsis *AtSPR1* overexpression vector under the control of CaMV 35S promoter and transformed it into the wild-type Col. The phenotype of the *AtSPR1* overexpressing transgenic plants was similar to that of *SmSPR1* overexpressing transgenic plants. In the dark, the hypocotyls of the etiolated seedlings also showed a right-handed helix phenotype after three days (Fig. 5D). In the presence of light, there was no difference between the transgenic plants and the wild-type phenotype (Supplemental Fig. S6). Semi-quantitative RT-PCR analyses revealed the transcription levels of the *AtSPR1* genes, which indicated that the right-handed helical phenotype was not the result of posttranscriptional gene silencing (Fig. 5E).

### SPR1, CSN5A, and HY5 Physically Interact with Each Other *In Vivo*

The appearance of the right-handed helical orientation in etiolated but not in light-grown seedling prompted us to hypothesize that the helical phenotype is related to light. We then transferred the helical etiolated seedling (5 days) to light conditions, which resulted in straight, newly grown upper hypocotyls (stems) after 4 days of growth, and the formed lower stems also showed right-handed helical orientation (Fig. 6A). On the other side, we transferred the light-grown seedling (5 days) to the dark environment, and after 6 days of growth, the upper hypocotyl cells rapidly elongated and formed right-handed helical epidermal cells (Fig. 6B). Taken together, our data demonstrated that the right-handed helical orientation of upper epidermal cells in transgenic plants was directly triggered by light. Therefore, we hypothesized that there are light-regulated proteins that can bind to SPR1 and participate in anisotropic cell growth. To verify this hypothesis, we next screened the interaction protein of SPR1 using a pull-down assay, yeast two-hybrid (Y2H), and a biomolecular fluorescence complementation (BiFC) assay.

**Fig. 6.**
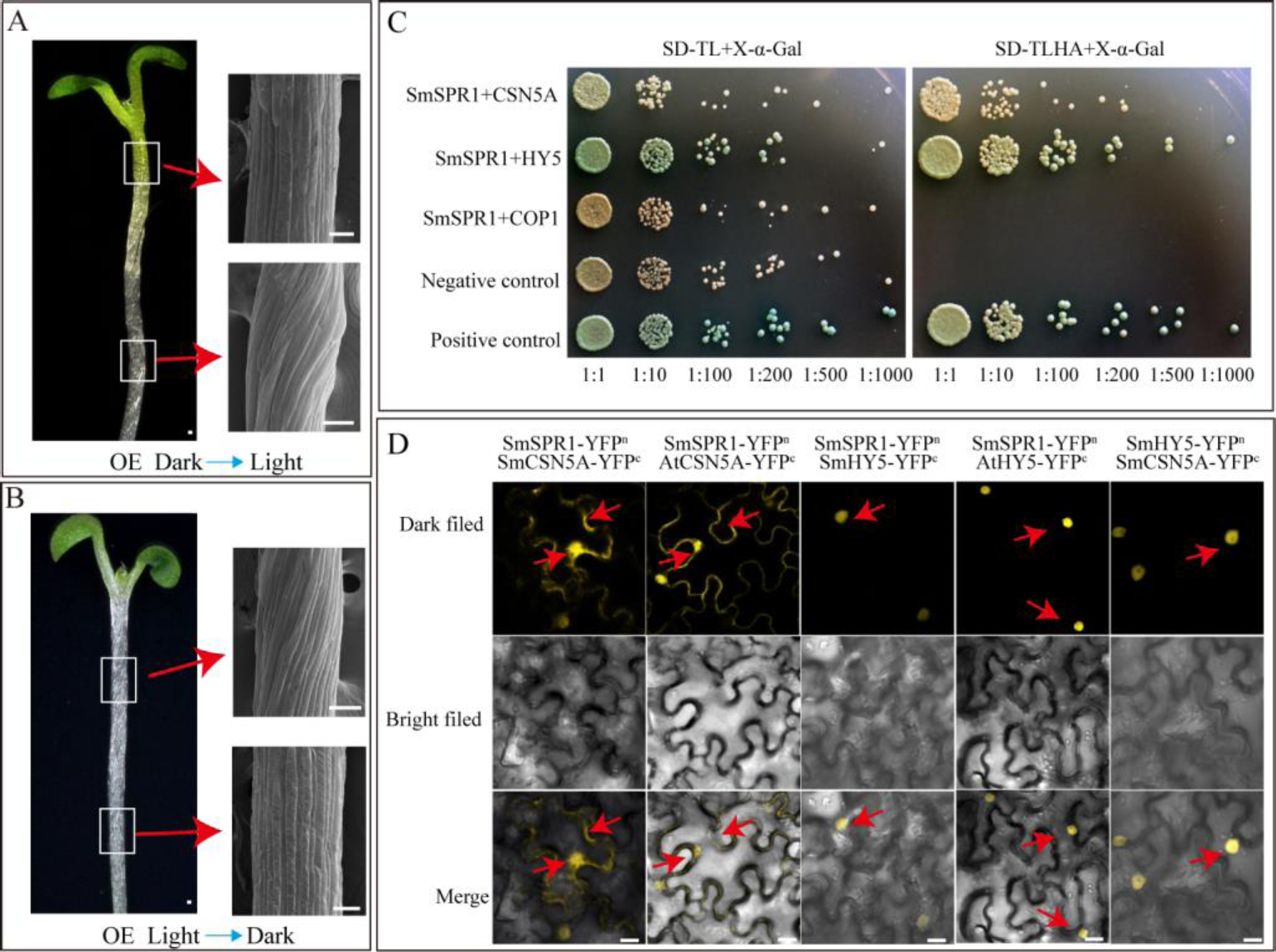
SmSPR1, CSN5A, and HY5 interaction in vivo. (A) Five-day-old etiolated seedlings were transferred to light conditions for another four days of growth. Bars = 10 µm. (B) Five-day-old light-grown seedlings were transferred to the dark for another six days of growth; Red arrows show scanning electron micrographs of the hypocotyl. (C) Interactions of SPR1 and CSN5A, HY5, and COP1 in a Y2H system. (D) SPR1, CSN5A, and HY5 interact with each other in the BiFC system. Bars= 10 µm.

Pull-down assays of plant extracts were conducted to identify proteins that physically interact with SPR1. Glutathione S-transferase (GST)-fused SPR1 proteins were first expressed in an *Escherichia coli* system and affinity purified. Recombinant GST alone and GST-SPR1 were then used as baits to identify the interacting proteins. SDS-PAGE gels were used to separate precipitates, which were then analyzed by matrix-assisted laser desorption ionization time-of-flight mass spectroscopy (MALDI-TOF-MS). A total of 1,145 peptide fragments that could be assembled into 451 proteins were identified, and different proteins detected between GST alone and GST-SPR1 were used as baits for functional annotation in the Nr database. We then conducted a yeast two-hybrid screen to detect the interacting proteins of SmSPR1, and a total of 11 proteins were identified. By combining these two results, three light-related proteins (CSN5A, HY5, and COP1) were next selected for further validation. We conducted a yeast two-hybrid one-to-one verification, which indicated that CSN5A and HY5, but not COP1, interacted with SPR1 (Fig. 6C). To confirm whether these three proteins could interact with SPR1 *in vivo*, we performed a BiFC assay using tobacco epidermal cells in both dark and light conditions. In the presence of light, co-expression of SPR1-CSN5A showed strong YFP fluorescence in the nuclei and cytomembrane, whereas SPR1-HY5 was only co-expressed in the nuclei (Fig. 6D), and no fluorescence was observed in the SPR1-COP1 combination. In the dark, co-expression of SPR1-CSN5A also exhibited strong YFP fluorescence, whereas the SPR1-HY5 combination was not reconstituted in the functional YFP (Supplemental Fig. S7). The fact that both CSN5A and HY5 physically interact with SPR1 prompted us to investigate whether CSN5A and HY5 interact with each other using BiFC, which revealed a strong physical interaction in the presence of light (Fig. 6D).

## DISCUSSION

Amino acid sequence alignment of SmSPR1s showed that, similar to the Arabidopsis SPR1 family proteins, the amino acid sequence of SmSPR1s is conserved at the N- and C-termini (Fig. 1B). Previous studies have demonstrated that the N- and C-termini of AtSPR1s can bind to MTs individually and perform partial functions (Nakajima *et al*., 2004). Therefore, we speculate that the N- and C-termini of SmSPR1s also have core functions, and the middle region with a certain length and a random sequence is only used as a connection and spacer between the two core regions. Interestingly, SmSPR1_L5 is conserved only at the N-terminus of the amino acid, whereas the C-terminus is a variable region. Combined with the results of real-time PCR, the expression level of the SmSPR1_L5 gene is extremely low in all assayed tissues (Fig. 1C). We inferred that *SmSPR1_L5* is a redundant gene, and it is possible that its function has been lost.

A previous study has shown that the root epidermal cells of overexpression Arabidopsis AtSPR1 transgenic lines are indistinguishable from the wild type, and increased *AtSPR1* expression does not cause root twisting. The cell elongation kinetics of the *P35S:AtSPR1* transgenic lines and the wild type were also compared using dark-grown hypocotyls, while in this process, the helical phenotype was also not observed (Nakajima *et al*., 2004; Sedbrook *et al*., 2004). However, a right-handed helical orientation of hypocotyl epidermal cells in dark-grown seedlings was observed in the *P35S*:*SmSPR1* transgenic lines (Figs. 2A, B), which raised the question of whether overexpression of Arabidopsis AtSPR1 results in a helical phenotype. To address this, overexpression of Arabidopsis *AtSPR1* was repeated, which showed a similar right-handed helix in etiolated seedlings compared to *SmSPR1* overexpressing transgenic plants (Fig. 5D). An Arabidopsis *spr1* mutant rescue experiment was also conducted, and the *P35S:SmSPR1* overexpression lines were rescued from the oblique phenotype involving the roots and helical growth of epidermal cell files (Figs. 5A, B). Taken together, previous studies have indeed neglected the helix phenotype of etiolated seedlings in overexpression *AtSPR1* transgenic lines, and we inferred that the SmSPR1 and AtSPR1 generally have a similar function of regulating plant phenotypes.

In the dark, etiolated seedlings enter a rapid elongation period after three days, and the elongation zone moves up from the base-middle regions to the apical third of the hypocotyl (Gendreau *et al*., 1997). Coincidentally, the helix phenotype of *SPR1*-overexpressing transgenic plants is expressed only at the upper zone of hypocotyls of etiolated seedlings after three days, which is exactly the time and zone at which the etiolated seedlings grow rapidly (Fig. 2D). Therefore, the helical phenotype may be related to the rapid growth of the hypocotyl in the dark.

In the Arabidopsis *spr1* mutant, three major phenotypes differed from the wild type, which include roots undergoing oblique growth to the right, helical growth of hypocotyls in both light- and dark-grown seedlings, and hypocotyl helices expanding and shorting in etiolated seedlings. However, in our *P35S:SmSPR1* transgenic lines, only the phenotypes of right-handed helix and expansion of etiolated seedling were observed, which prompted us to find light-correlated factors that interact with SPR1 to regulate cell expansion and seedling growth. Finally, two light-correlated factors, CSN5A and HY5, were identified and confirmed to interact with SPR1 *in vivo*. The CSN (COP9 signalosome), which was identified as a photomorphosis inhibitor in Arabidopsis, consists of eight subunits (Wei *et al*., 1994). CSN is a metalloproteinase that cleaves covalently linked RUB1/NEDD8 from CRL E3 ubiquitin ligase of cullin proteins, and this process is known as derubylation/deneddylation (Lyapina *et al*., 2001). CSN5 is one of the core subunits in the COP9 signalosome, and the derubylase catalytic center is located at the conserved JAMM motif of the CSN5 subunit (Cope *et al*., 2002). Previous studies have shown that SPR1 can be ubiquitinated under salt stress and then degraded by the 26S proteasome (Wang *et al*., 2011). The interaction between SPR1 and CNS5A indicates that the COP9 signalosome participates in SPR1 ubiquitination. CSN also has kinase activity or can bind to kinase-associated proteins and may exist in post-translational modification mechanisms (such as phosphorylation) (Meister *et al*., 2016). MT end-binding protein EB1 can degraded by the ubiquitin system, and CSN has the ability to stabilize EB by binding to it via the CSN5 subunit, which mediates phosphorylation of EB1 and prevents EB1 from degradation (Peth *et al*., 2007). Whether SPR1 can also be phosphorylated by CSN needs further investigation.

HY5 is a member of bZIP transcription factor family that regulates fundamental developmental processes such as inhibition of hypocotyl growth, lateral root development, cell elongation, pigment accumulation, and other phenotypes related to photomorphogenesis (Gangappa and Botto, 2016). In the dark, HY5 can be degraded by ubiquitination of COP1, thereby inhibiting the occurrence of light morphology (Oyama *et al*., 1997). Keech et al. proposed that HY5 is repressed in destabilizing the cortical MT array in the epidermis and mesophyll cells during leaf senescence in the dark (Olivier *et al*., 2011), i.e., HY5 can indirectly promote the stability of MTs, which is similar to the function of SPR1. Our Y2H and BiFC assays showed that SPR1 interacts with HY5 *in vivo*, which prompted us to hypothesize that SPR1 and HY5 synergistically facilitate MT stabilization in the presence of light. HY5 also positively controls cell proliferation in the secondary thickening and negatively regulates lateral root formation, which may influence cell mitosis and division (Chen and Han, 2016; Oyama *et al*., 1997). Taking this together with the fact that MTs are involved in cell division as major components of spindle and SPR1 interacts with HY5, we propose that SPR1 and HY5 interact with each other during the cell cycle. To conclude, in view of these new findings, we propose a tentative model wherein SPR1 regulates the morphology of MTs by interacting with different proteins in light and dark conditions. In the presence of light, SPR1 binds to CSN and HY5. In the dark, SPR1 only binds to CSN to regulate the morphology of MTs, further regulating cell elongation and directional organ growth (Fig. 7A).

**Fig. 7.**
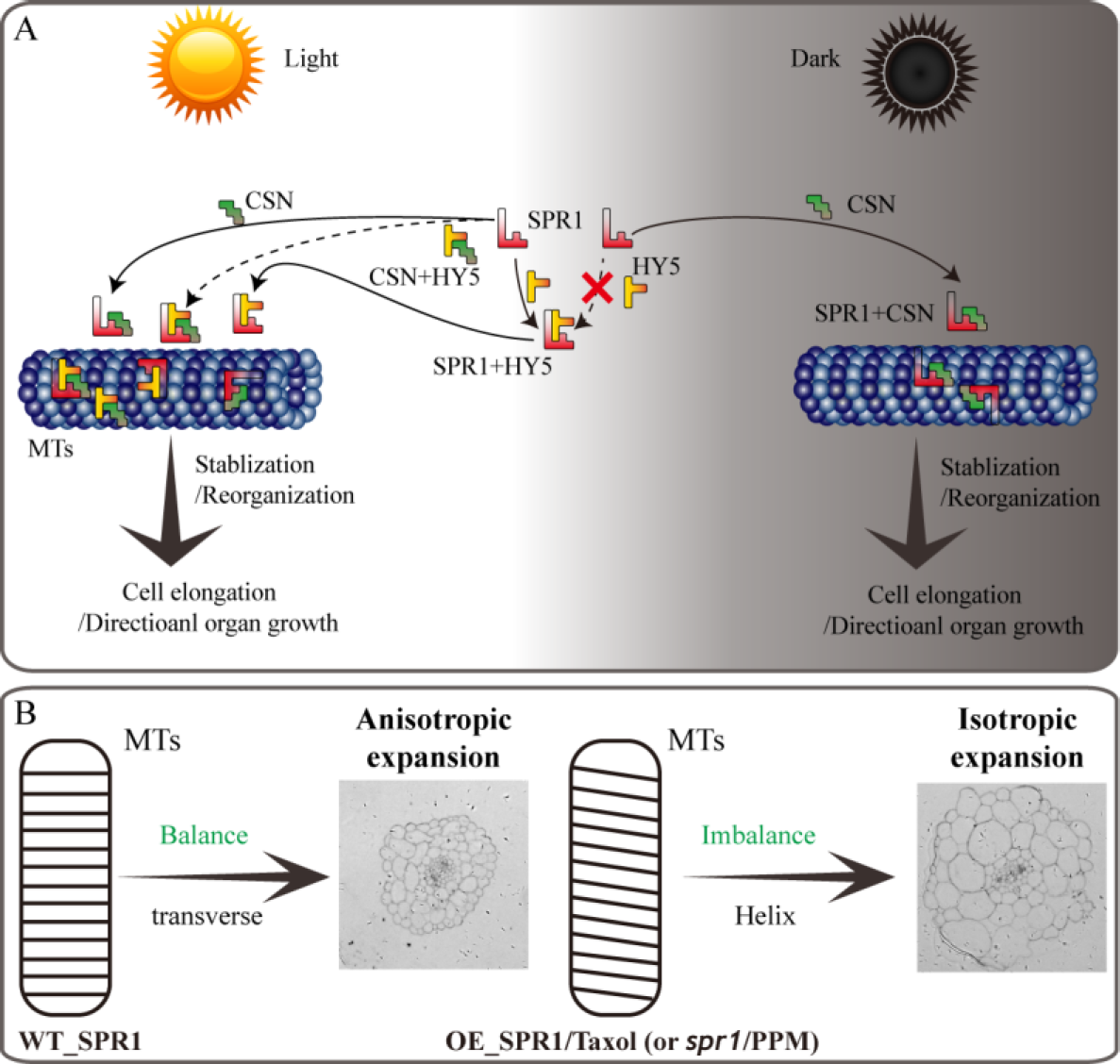
Model of SPR1, CSN, and HY5 interactions and helical growth. (A) In the light, SPR1, CSN, and HY5 could interact to each other for the stabilization or reorganization MTs and ultimately mediate cell elongation or directional organ growth. In the dark, SPR1 only interacts with CSN to control the next MT-related biological process. (B) In wild-type plants, the expression level of SPR1 accords with the balance of stability and polymerization of MTs. MTs arrays are transverse to the elongation axis, which result in a normal anisotropic expansion of cells. Meanwhile, in the SPR1 overexpression transgenic, *spr1* mutant, and MTs drug treatments plants, the balance of MTs was disrupted, which in turn led to the helix of the MTs and following isotropic expansion of cells.

The reason for the helix of *spr1* transgenic plants is caused by isotropic cell expansion, which there is widespread consensus about. However, the cause of the isotropic expansion of cells has been controversial, particularly with regard to whether the isotropic growth of hypocotyl cells is caused by changes in the morphology of microtubule arrays. Furutani et al. (2000) suggested that a defect in MT organization results in reduced anisotropic growth. MT arrays are more irregularly oriented in ground tissue cells than in epidermal cells in etiolated *spr1* hypocotyls, and this varying degree of MT defects cause isotropic cell growth(Furutani *et al*., 2000). Nakajima et al. (2004) also observed oblique MT arrays in *spr1* hypocotyls and roots, and they agree that defective MTs cause isotropic cell growth that further leads to a helix morphology of tissues (Nakajima *et al*., 2004). However, Sedbrook et al. (2004) obtained different results in relation to the *spr1* phenotype; they found that directional cell expansion is discordant with MT oblique arrangements and suggested that the pitch of MTs in epidermal cells does not guide epidermal cell file twisting (Sedbrook *et al*., 2004). Our study found that hypocotyls of etiolated seedlings in *SPR1*-overexpressing transgenic plants showed a pronounced phenomenon of oblique MT arrays. However, at roots where helix growth does not occur, the morphology of MTs did not exhibit any changes, and there was no difference in the morphology of MTs between the transgenic and wild-type plants in the presence of light (Figs. 3E, F). Thus, we suggest that the oblique MT array is directly related to isotropic cell growth. However, whether the oblique MT array leads to isotropic cell growth or isotropic growth of cells causes obique of MTs could not be determined in the present study. We prefer that SPR1 affects the morphology of MTs, which leads to a decrease in anisotropic growth of cells. Because, the known function of SPR1 is to stabilize MTs that are related to the synthesis of cellulose, which is synthesized along MTs, and the arrangement of fibers is related to the morphology of the cell walls and cells (Himmelspach *et al*., 2003). Changes in MT structure could result in alterations in the alignment of cellulose, which in turn leads to modifications in cell morphology.

Furutani et al. (2000) proposed a model for the *spr1* helical phenotype. In the axial direction of cell tissues, loss of anisotropic growth of inner cells leads to cell shortening, which ultimately results in plant shortening. Compared to shortened internal cells, the length of epidermal cells does not change. To compensate for this difference, epidermal cells are skewed to maintain the same length as internal cells(Furutani *et al*., 2000). This model has been used to explain the spiral phenotype of *spr1* (Nakajima *et al*., 2004; Nakajima *et al*., 2006; Sedbrook *et al*., 2004). Overexpression of *SmSPR1* was also observed in the helix phenotype; the results of semi-thin section assays and PI staining experiments showed isotropic growth and expansion of cells in transgenic plants (Figs. 3B, D). However, the hypocotyls of *SmSPR1* overexpressing transgenic lines did not become shorter (Fig. 2). The shortening of hypocotyls is one of the main characteristics of the above model, which is discordant with our results and cannot explain the *SmSPR1* overexpression helix phenotype. According to the model, *SmSPR1*-overexpressing transgenic lines both have helixes, and thus, the absence of shortened hypocotyls is a contradictory finding. How do we explain this phenotype of *SmSPR1* overexpressing transgenic lines? Nakajima et al. (2004) found that the overexpression of *AtSPR1* causes a small increase in the elongation of hypocotyls (Nakajima *et al*., 2004). Taken together, our results suggest that the contradictory phenotype of *SmSPR1* overexpression transgenic lines could be attributable to both increased elongation and isotropic growth of hypocotyl cells. We can also assume that the expression level of *SPR1* in the Arabidopsis wild type does not meet the maximum elongation of the hypocotyl, and increased expression of *SPR1* may accelerate hypocotyl elongation.

How can we explain the spiral phenomenon of hypocotyl caused by overexpression of SPR1? We propose a model in which the SPR1 regulates MT polymerization in addition to stabilizing MT. The stability and polymerization of MTs determines the morphology of MTs. Both the *spr1* mutant and overexpressing *SPR1* lines showed a disruption of this balance, which in turn lead to the helical orientation of the MTs (Fig. 7B). Our pharmacology experiments data demonstrated that *SmSPR1* overexpressing transgenic plants have strong MT depolymerization tolerance relative to the wild-type plants (Figs. 4C-G), demonstrating that MTs of transgenic plants are more stable than those of wild-type plants. Taxol (paclitaxel) is an MT-stabilizing drug often used for MT stability studies. Taxol and propyzamide both caused identical handedness of Arabidopsis seedlings, despite they have the opposite effect on the polymerization of MTs (Furutani *et al*., 2000; Sedbrook *et al*., 2004). In wild-type seedlings, 0.3 µM taxol leading to a clockwise spiral of cotyledons, and 1 µM taxol resulting in root reduced growth and left-hand bending (Furutani *et al*., 2000). Both SPR1 and taxol act to stabilize the MTs; increased expression of *SPR1* and adding taxol both result in the helix phenotype of Arabidopsis, which indirectly demonstrates that overexpression of SPR1, like the addition of taxol, can cause MTs to be over stabilized, thus resulting in changes in the morphology of MTs and plants.

## Acknowledgments

This work was supported by The Fundamental Research Funds for the Central Non-profit Research Institution of Chinese Academy of Forestry (CAFYBB2018QB001). We thank LetPub (www.letpub.com) for its linguistic assistance during the preparation of this manuscript.

## Supplementary data

Supplementary data are available at JXB online.

**Table S1**. List of primers used for cloning *SmSPR1, SmCSN5A, SmHY5* and *SmCOP1*

**Table S2**. List of primers used for Semi-quantitative RT-PCR of *SmSPR1* and *AtSPR1*

**Table S3**. List of primers used for real-time RT-PCR analysis

**Table S4**. List of primers used for overexpression conduct SmSPR1 and AtSPR1

**Table S5**. List of primers used for construction of prokaryotic expression vector

**Table S6**. List of primers used for Yeast Two-Hybrid

**Table S7**. List of primers used for BiFC.

**Fig. S1**. Phenotype of overexpression SmSPR1 in the condition of light. (A) Seedling phenotype between wild-type and *SmSPR1* transgenic plants. (B) Stem cross section of wild-type and transgenic Arabidopsis plants. 1-5 show five equal parts from shoot tip of the stem to the base. (C) Micrographs of the upper and lower hypocotyl and root. (D) PI stain of hypocotyl and root of the wild-type and transgenic seedlings in the presence of light. Bars = 25 µm.

**Fig. S2**. PI staining of hypocotyl and root of the wild-type and transgenic etiolated seedlings from three to seven days of growth. Bars = 100 µm.

**Fig. S3**. Hypocotyl growth trajectory of wild-type and transgenic etiolated seedlings from three to seven days of growth. Data are expressed as the mean ± SD of > 30 seedlings. Significant differences were determined using the Student’s *t* test (*P* < 0.01).

**Fig. S4**. The SmSPR1: GFP localization. (A) Confocal images of SmSPR1: GFP (green). (B) PI-stained root (red) in the condition of light. Bars = 20 µm.

**Fig. S5**. SmSPR1 transgenic seedlings have increased PPM tolerance. (A) Seedling phenotypes of the wild-type and transgenic plants on culture medium containing PPM. (B) PI staining of hypocotyl and roots of the wild-type and transgenic plants with the increasing concentrations of PPM in the dark. Bars = 50 µm.

**Fig. S6**. Seedling phenotype in the wild-type and *AtSPR1* transgenic plants. Bars = 1 cm.

**Fig. S7**. Interactions among SPR1, COP1, and HY5 using the BiFC system. (A) Negative control. (B) Positive control. (C) SmSPR1 interact with CSN5A and HY5 in the dark. (D) SmSPR1 interact with COP1 in both light and dark conditions. Bars = 20 µm.

